# Temporal niche expansion in mammals from a nocturnal ancestor after dinosaur extinction

**DOI:** 10.1101/123273

**Authors:** Roi Maor, Tamar Dayan, Henry Ferguson-Gow, Kate E. Jones

## Abstract

Most modern mammals, including strictly diurnal species, exhibit sensory adaptations to nocturnal activity, thought to be the result of a prolonged nocturnal phase or ‘bottleneck’ during early mammalian evolution. Nocturnality may have allowed mammals to avoid antagonistic interactions with diurnal dinosaurs during the Mesozoic. However, understanding the evolution of mammalian activity patterns is hindered by scant and ambiguous fossil evidence. While ancestral reconstructions of behavioural traits from extant species have the potential to elucidate these patterns, existing studies have been limited in taxonomic scope. Here, we use an extensive behavioural dataset for 2415 species from all extant orders to reconstruct ancestral activity patterns across Mammalia. We find strong support for the nocturnal origin of mammals and the Cenozoic appearance of diurnality, although cathemerality (mixed diel periodicity) may have appeared in the late Cretaceous. Simian primates are among the earliest mammals to exhibit strict diurnal activity, some 52-33Mya. Our study is consistent with the hypothesis that temporal partitioning between early mammals and dinosaurs during the Mesozoic led to a mammalian nocturnal bottleneck, but also demonstrates the need for improved phylogenetic estimates for Mammalia.

---

Although mammals exhibit striking morphological, behavioural and ecological niche diversity^1^, the distribution of mammalian activity patterns is strongly biased towards nocturnality^2^ Additionally, most mammalian species, including strictly diurnal ones, exhibit visual adaptations to nocturnal activity that are similar to those of nocturnal birds and reptiles^3^. For example, mammals lack photoreception mechanisms (e.g. parietal organs) that are found in other amniotes ^3^, and exhibit reduced diversity of active photoreceptors^4,5^. Additionally, there is evidence that enhanced olfactory sensitivity^6^, broader frequency range hearing^7^, and sophisticated whisker-mediated tactile perception^8^ may have evolved in mammals to compensate for insufficient visual information^3,5^. In general, species exhibit characteristic patterns of activity distribution over the 24-hour (diel) cycle, and as environmental conditions may change radically, yet predictably between day and night, activity patterns allow individuals to anticipate fluctuations, and time activity optimally^9,10^. Physiological and behavioural adaptations to different activity patterns are significant to individual fitness^11^, and therefore to species evolutionary success^12,13^. Moreover, long-term shifts in activity patterns may reveal shifts in selective regimes, caused by changes in biotic and abiotic conditions^13,14^.

The predominance of nocturnal adaptations in mammals may be the result of a prolonged nocturnal phase in the early stages of mammalian evolution, after which emerged the more diverse patterns observed today^5,15^. This ‘nocturnal bottleneck’ hypothesis suggests that mammals were restricted to nocturnal activity by antagonistic interactions with the ecologically dominant diurnal dinosaurs during the Mesozoic^5,15,16^. The Cretaceous-Paleogene (K-Pg) mass extinction event circa 66Mya, led to the extinction of all non-avian dinosaurs along with the marine- and flying reptiles, and the majority of other vertebrates, and invertebrate and plant taxa^17,18^. This event marks the end of the Mesozoic ‘reign of dinosaurs’ and the transition to the mammal-dominated Cenozoic fauna. If an antagonistic interaction with dinosaurs was an important factor in restricting early mammals to nocturnal activity, then all Mesozoic mammals are expected to have been nocturnal, and diurnal mammals would have only appeared after the K-Pg mass extinction event.

Support for the nocturnal bottleneck hypothesis remains indirect. For example, some Synapsids, the non-mammalian lineage ancestral to mammals, were adapted to nocturnal activity >300Mya, suggesting a nocturnal origin for mammals^19^. However, as all modern mammals (except monkeys and apes) have nocturnal morphological adaptations regardless of their activity^3,20^, inferring activity patterns from fossil cranial morphology may be unreliable. Evidence from genetic and histological studies of the evolutionary development of mammalian eyes indicate that nocturnal adaptations preceded diurnal ones^4,21^, but this does not help elucidate questions around the timing of these adaptations. Ancestral reconstructions of behavioural traits using a phylogenetic comparative approach may help to understand both the pattern and timing of the evolution of activity patterns in mammals since activity patterns have been shown to be genetically determined^22^ yet responsive to selective pressures^10^. However, phylogenetic studies of mammalian activity patterns so far have focused on two mammalian orders – primates^23-25^ and rodents^26^. Primate activity patterns have been studied extensively, and some evidence suggests that primate diurnality originated in the most recent common ancestor (MRCA) of suborder Haplorrhini (all monkeys, apes and tarsiers)^13^ in the Mesozoic^27,28^. It is conceivable, although thus far not tested, that diurnal diversifications in other orders of Mesozoic origins, e.g. Scandentia (treeshrews), Macroscelidea (elephant shrews) and Rodentia, could have occurred before the extinction of dinosaurs, calling for a wider examination of how activity patterns evolved across mammals.

Here, we use an extensive dataset of activity patterns for 2415 mammal species, representing 135 of the 148 extant families and all extant orders (Supplementary Table 1) to investigate ancestral activity patterns in mammals, and to understand the timings of the appearance of mammal diurnality. We assign species to one of five activity patterns: (*i*) nocturnal – active only or mostly in the dark; (*ii*) diurnal – active only or mostly during daylight hours; (*iii*) cathemeral – active both during the day and during the night; (*iv*) crepuscular – active only at twilight, around sunrise and/or sunset; and (*v*) ultradian – active in cycles of a few hours. (see Methods). We map the three main activity patterns (nocturnal, cathemeral, and diurnal) onto two phylogenetic frameworks representing two of the main hypotheses of mammalian evolutionary history for our analyses, termed here short-fuse (SF) following^28^ updated by^29^, and long-fuse (LF) phylogenies (adapted from^27^) (Fig. 1). We then use reversible jump Markov Chain Monte Carlo (rjMCMC) methods^30^ to estimate transition rates between different activity states, and to infer the posterior probability (PP) of character states at each node in the phylogenies to examine the evolution of activity patterns of mammals and estimate support for the nocturnal bottleneck hypothesis.

**Figure 1.**
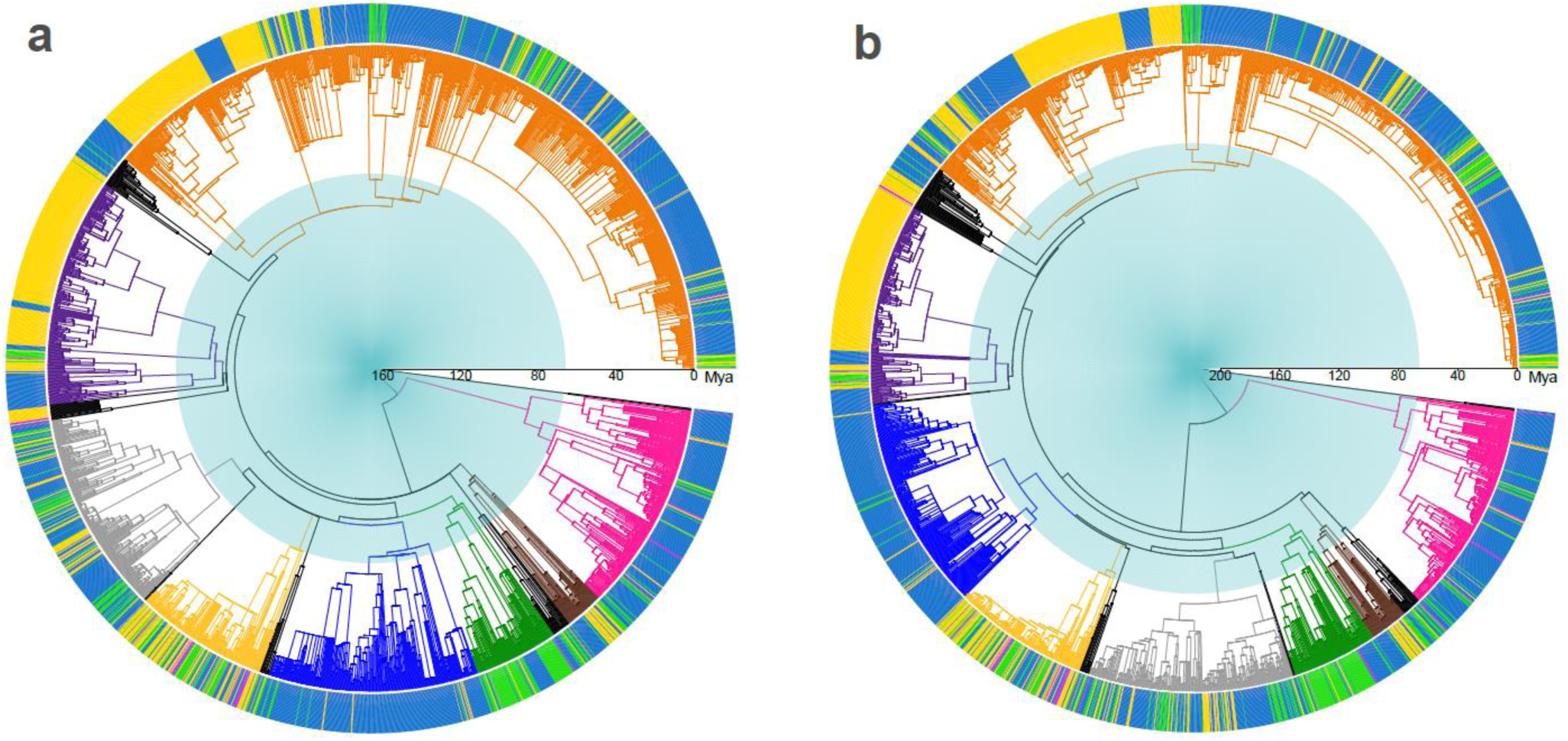
Activity patterns distribution across (a) the short-fuse (SF), and (b) the long-fuse (LF) estimates of mammalian evolution. Species activity patterns are denoted by different colours in the perimeter circle, where nocturnal is denoted as blue; diurnal yellow; cathemeral green; and ambiguous magenta. Branch colours represent taxonomy, where Marsupials are pink; Afrotheria brown; Soricomorpha+Erinaceomorpha green; Chiroptera blue; Cetartiodactyla yellow; Carnivora grey; Primates purple; Rodentia orange; and all other orders are black. Mesozoic and Cenozoic eras are denoted by blue and white backgrounds, respectively. SF phylogeny follows^28^ updated by^29^, and LF phylogeny is adapted from^27^ (see Methods). Branch lengths are proportional to time (Myr).

## Results

We find that the model values of PP_Noct_ (posterior probability of nocturnality) at the ancestral node of extant mammals were 0.74 (Credible Interval, CI 0.71-0.76) and 0.59 (CI 0.54-0.64) for SF and LF phylogenies, respectively, offering strong support for a noctural ancestor (Fig. 2). In contrast, a cathemeral or a diurnal ancestral state is less well supported: modal value of PP_Cath_ (posterior probability of cathemerality) = 0.24 (CI 0.23-0.26) and 0.31 (CI 0.29-0.33) for SF and LF, respectively, or PPDiur (posterior probability of diurnality) = 0.02 (CI 0.01-0.03) SF and 0.1 (CI 0.07-0.14) LF (Fig. 2). The narrow and non-overlapping distributions of PP values across the activity pattern reconstructions indicate that our results are consistent and robust across samples of the rjMCMC chains, although the distributions are wider using the LF phylogeny (Fig. 2).

**Figure 2.**
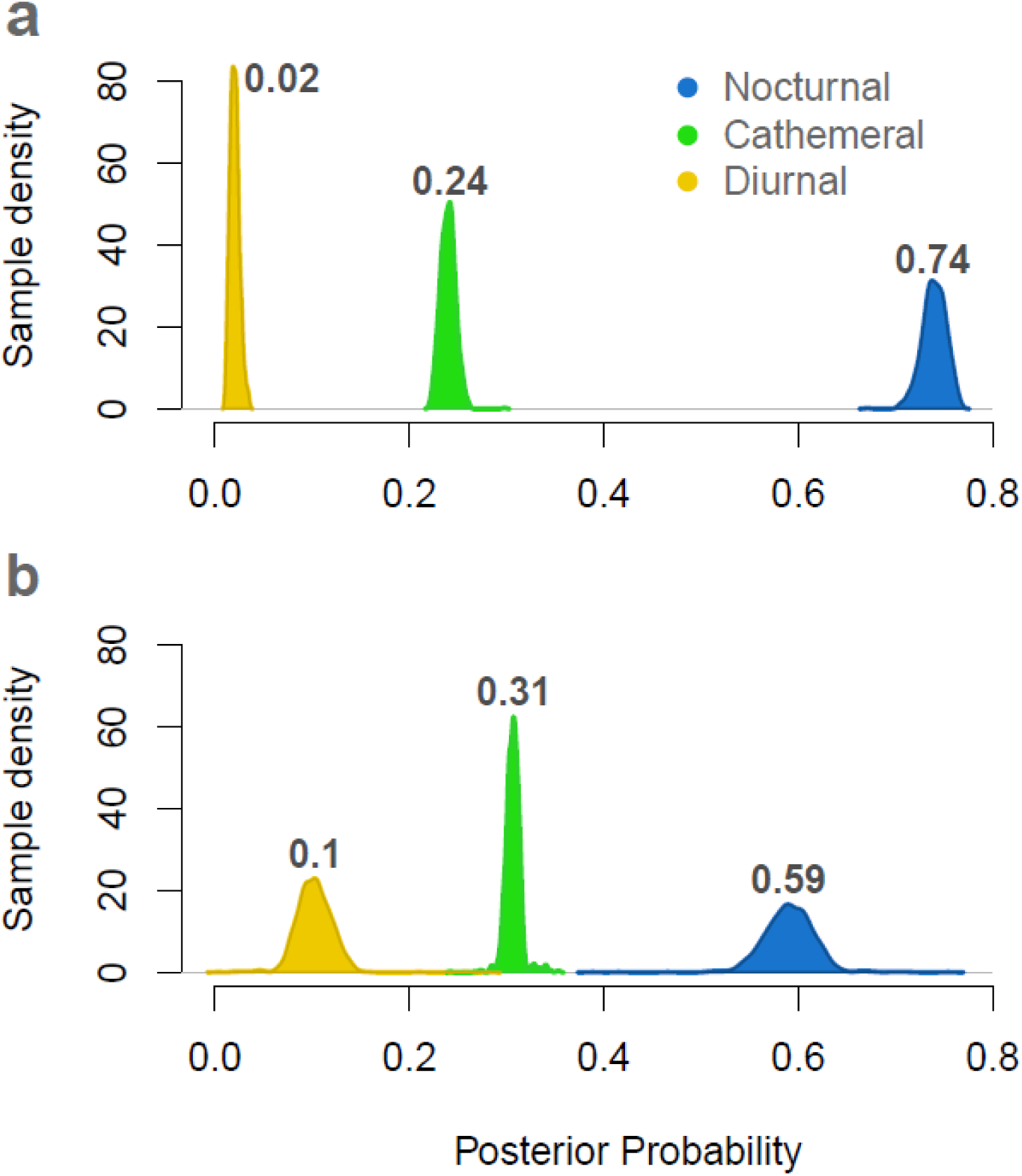
Posterior probability (PP) density of ancestral activity patterns reconstructions of the most recent common ancestor of crown-group Mammalia from (a) SF and (b) LF phylogenies. Distribution curves are calculated from 1000 post-burnin rjMCMC samples, and modal PP values for each distribution are shown in bold. Colours correspond to activity patterns.

The first strong evidence (where the reconstructed activity pattern was supported by modal PP values >0.67) in mammals of an expansion of temporal niche into cathemerality, is in the early Paleogene (Cenozoic) for the SF phylogeny (no later than 65.8Mya), or in the late Cretaceous (Mesozoic) for the LF phylogeny (no later than 74.7Mya) (Figs. 3 and 4). Although the LF phylogeny supports a Mesozoic shift to cathemerality, the modal PP values of the remaining 41 Mesozoic nodes were either nocturnal (23 nodes), or unclear – where all three activity patterns were supported by modal PP values <0.67 (18 nodes). Using the SF phylogeny, we reconstruct the first transition to cathemerality in the MRCA of order Cetartiodactyla (cetaceans and even-toed ungulates). This taxa was likely to be cathemeral (PP_Cath_ = 0.79 CI 0.72-0.87), and almost certainly exhibited considerable daytime activity (PP_Noct_ = 0.02 CI 0.01-0.04) (Fig. 3). Using the LF phylogeny, the first cathemeral transition was in the MRCA of families Soricidae (shrews) and Talpidae (moles) (PP_Cath_ = 0.81 CI 0.61-0.91; PP_Diur_ = 0.07 CI 0.03-0.15) (Fig. 4).

**Figure 3.**
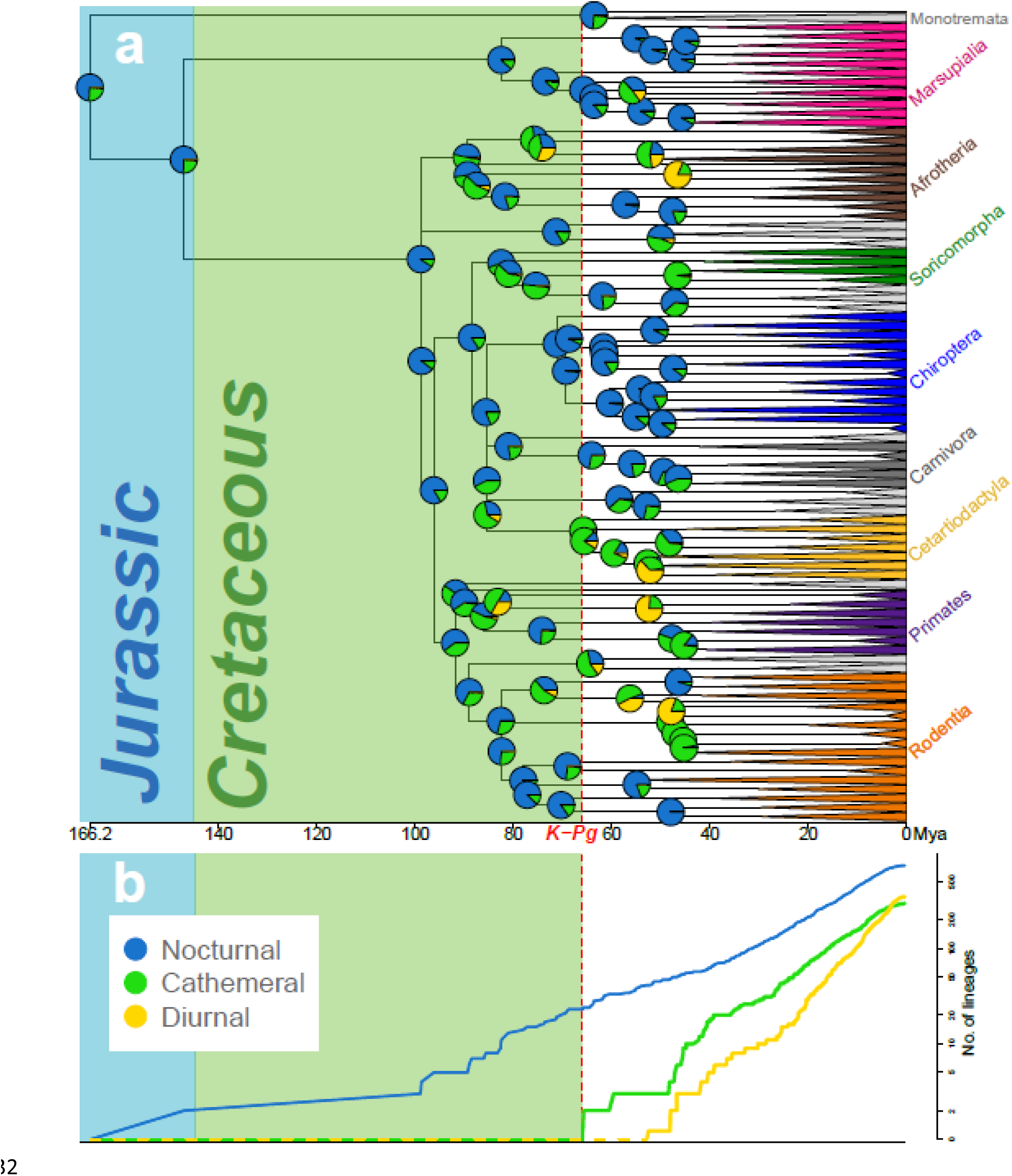
Reconstruction of ancestral activity patterns and character accumulation, across the ‘short fuse’ (SF) hypothesis of mammalian evolution. (a) Ancestral activity pattern reconstruction across the SF phylogeny^28^ updated by^29^. Pie charts correspond to ancestral reconstructions at each node, and colours denote the proportional value of the posterior probability (PP) of each activity pattern, where nocturnal is blue; cathemeral green; and diurnal yellow. Shading denotes geological era. Branch lengths are proportional to time, with branches younger than 45Mya replaced with wedges for visualisation purposes. The red dashed line represents the K-Pg boundary. (b) Lineages through time plot for activity patterns. The predominant activity pattern was assigned to each node based on PP values, with a minimum value of 0.67. Nodes with reconstructed activity pattern PP values of <0.67 were excluded from the lineages through time plot.

**Figure 4.**
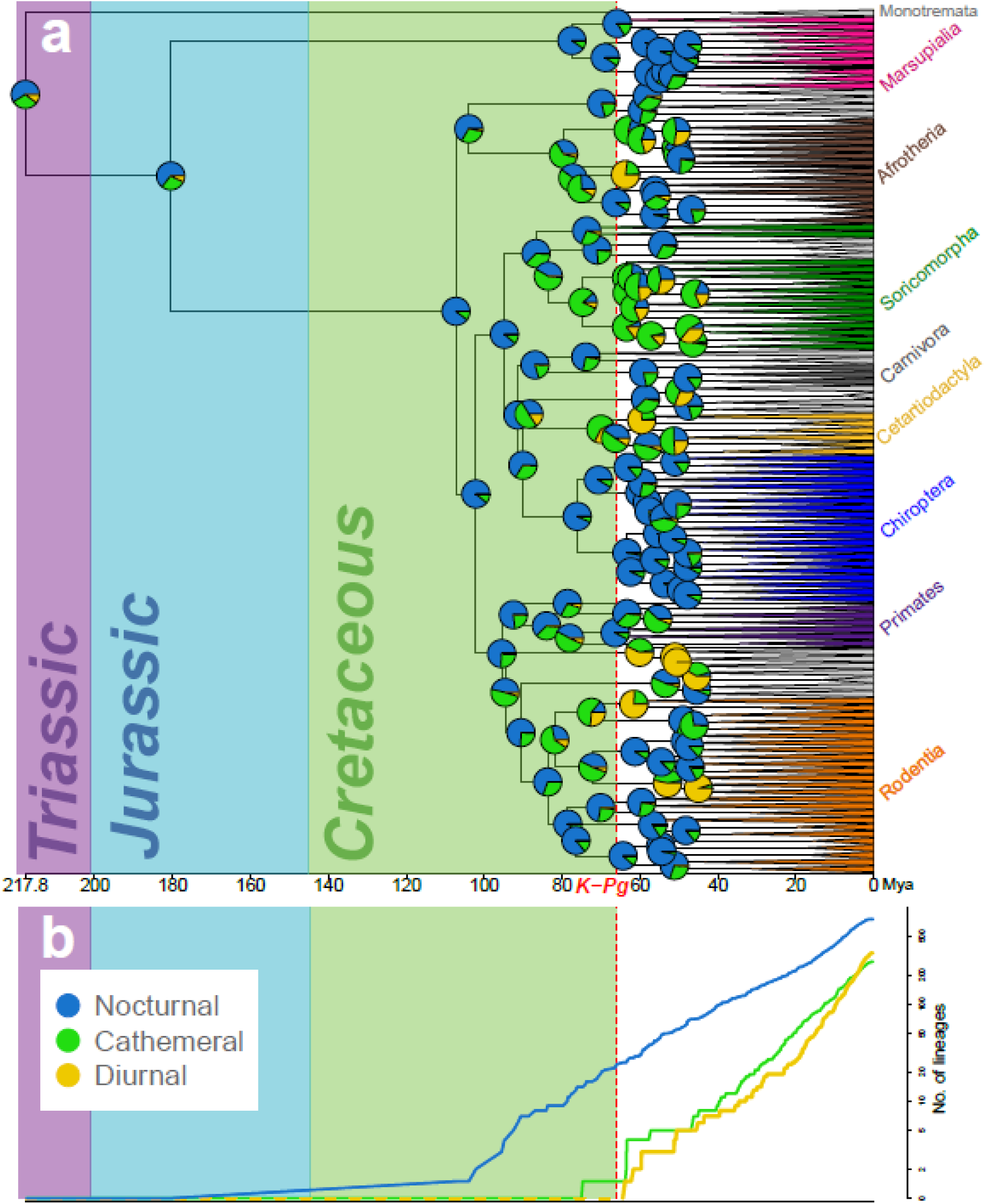
Reconstruction of ancestral activity patterns and character accumulation, across the ‘long fuse’ (LF) hypothesis of mammalian evolution. (a) Ancestral activity pattern reconstruction across the LF phylogeny adapted from^27^. Pie charts correspond to ancestral reconstructions at each node, and colours denote the proportional value of the posterior probability (PP) of each activity pattern, where nocturnal is blue; cathemeral green; and diurnal yellow. Shading denotes geological era. Branch lengths are proportional to time, with branches younger than 45Mya replaced with wedges for visualisation purposes. The red dashed line represents the K-Pg boundary. (b) Lineages through time plot for activity patterns. The predominant activity pattern was assigned to each node based on PP values, with a minimum value of 0.67. Nodes with reconstructed activity pattern PP values of <0.67 were excluded from the lineages through time plot.

Evidence of the evolution of diurnality (modal PP values >0.67) first appears in the early Paleogene (no later than 52.4Mya or 63.8Mya for SF and LF phylogeny, respectively) (Figs. 3 and 4). Using the SF phylogeny, we reconstruct transition to diurnality in the MRCA of the Simiiformes (all monkeys and apes) (PP_Diur_ = 0.76, CI 0.75-0.78; PP_Cath_ = 0.23, CI 0.22-0.25) (Fig.3). Using the LF phylogeny, the first taxon to exhibit diurnal activity was the MRCA of the family Macroscelididea (elephant shrews) (PP_Diur_ = 0.77, CI 0.76-0.80; PP_Cath_ = 0.22, CI 0.19-0.23; 63.8Mya), followed by the MRCA of families Ctenodactylidae (comb rats, Rodentia) (PP_Diur_ = 0.76; CI 0.73-0.78; 61.6Mya), Camelidae (Cetartiodactyla) (PP_Diur_ = 0.74, CI 0.72-0.77; 59.6Mya), and Tupaiidae (treeshrews, Scandentia) (PP_Diur_ = 0.99, CI 0.99-0.99; 51.1Mya) in rapid succession (Fig. 4).

For both SF and LF phylogenies, we find that transition rates from a cathemeral pattern to either noctural or diurnal are about three times higher than the transition rates from either nocturnal or diurnal to cathemeral (Table 1). Furthermore, the transition rates in the SF reconstruction are three orders of magnitude lower than the respective rates in the LF reconstruction.

**Table 1.** Character transition rate matrix for SF and LF ancestral activity pattern reconstructions. Transition rates are from the state in the column to the state in the row and represent model posterior values. Direct transitions between nocturnal and diurnal were not allowed (0) under our character state transition model.

| Phylogeny |  | Transition rates |  |  |
| --- | --- | --- | --- | --- |
|  |  | Nocturnal | Cathemeral | Diurnal |
| Short fuse | Nocturnal | - | 0.01 | 0 |
|  | Cathemeral | 0.03 | - | 0.03 |
|  | Diurnal | 0 | 0.01 | - |
|  |  | Nocturnal | Cathemeral | Diurnal |
| Long fuse | Nocturnal | - | 1.97 | 0 |
|  | Cathemeral | 7.46 | - | 7.41 |
|  | Diurnal | 0 | 1.96 | - |

## Discussion

We have shown that extant mammals likely originated from a nocturnal ancestor, and that these ancestors remained nocturnal throughout the Mesozoic until either 9 Myr before the K-Pg event (LF reconstruction), or just after it (SF reconstruction). On balance, our evidence suggests that mammals likely remained nocturnal throughout the Mesozoic as nocturnal activity is strongly supported at most Mesozoic nodes in both SF and LF reconstructions. We find strong evidence that the shift to strict diurnality occurred after the K-Pg event (both SF and LF reconstructions), although cathemerality may have appeared in the late Cretaceous (74.7Mya LF reconstruction). Combined with other sources of evidence^5,21^, our analysis helps to further establish the nocturnal ancestry of mammals and the timing of the evolution of diurnality, as predicted by the nocturnal bottleneck hypothesis, namely that diurnality only originated in mammals after the disappearence of the dinosaurs.

Even if we accept the appearance of cathemeral mammals as an expansion of the temporal niche before the K-Pg event, it does not necessarily provide strong evidence against the nocturnal bottleneck hypothesis. Declines in dinosaur diversity long before the K-Pg event have been suggested, either globally, starting at least 40Myr before the K-Pg event^31^, or locally – herbivorous dinosaurs in present-day North America were declining for up to 15Myr prior to the event^18^. In contrast, fossils show that mammals had evolved considerable eco-morphological diversity as early as the mid-Jurassic period (174-164 Mya), and diversified along all axes of the ecological niche^32,33^, except the temporal axis. Moreover, extensive mammal radiations occurred following the Cretaceous Terrestrial Revolution (KTR, 120-80Mya), whereby angiosperms rose to dominate the global flora, and revolutionised eco-space^27,34,35^. Under such conditions, an invasion of mammals into the temporal niche of declining dinosaurs does not violate the assumption of temporal partitioning.

The MRCA of infraorder Simiiformes (monkeys and apes) was among the first taxa to have evolved diurnality (52.4Mya, SF reconstruction), and this is consistent with their evolution of diurnally-adapted vision^3,4,20^ – a singularity in mammals. Other diurnal clades such as squirrels (Sciuridae) and elephant-shrews (Macroscelididea) evolved at about the same time as the Simiiformes^27,28^ and presumably had similar opportunity to evolve diurnally-adapted vision, suggesting that diurnality in Simiiformes may have evolved considerably earlier than the minimum date of 52.4Mya. Simiiformes lie on an evolutionary branch that originates 83.2Mya (SF), when they diverged from tarsiers – their closest living relatives in the suborder Haplorrhini. Tarsiers are strictly nocturnal, but share with the Simiiformes several adaptations for high visual acuity, typical to diurnal vision^25,36^. The morphological and physiological adaptations to nocturnality in tarsiers are unlike those of any other nocturnal primate, suggesting that tarsiers originated from a diurnal ancestor, the MRCA of Haplorrhini, and secondarily adapted to nocturnal life^13^. The Haplorrhine MRCA was a Mesozoic species that lived until 83.2Mya (SF) or 78.1Mya (LF). This would imply that Mesozoic mammals were able to break out of the nocturnal bottleneck and endure direct interaction with dinosaurs following the KTR. Nevertheless, both reconstructions here, as well as other reconstructions of primate activity patterns based on different sets of data, including data on visual physiology, find weak or no evidence to the diurnality of the Haplorrhine MRCA^23-25^.

There are other uncertainties around the dates for three of the four taxa identified as shifting to diurnality within 7Myr after the K-Pg in the LF reconstruction (Macroscelididea, Ctenodactylidae, Camelidae). This is due to how we re-scaled the terminal-branches in^27^ to produce the species-level LF phylogeny. However, according to the dates given in^27^ and additional studies supporting the LF hypothesis^37-40^, these families originated in the Cenozoic, so our prediction of Cenozoic origins to mammal diurnality remains intact. The MRCA of Tupaiidae (Scandentia) and their closest living relative – the nocturnal Ptilocercidae (Pen-tailed tree shrews, a monotypic family) - has been placed in the Cenozoic, 60.1 Mya^27^ The LF reconstruction shows that this species was probably diurnal or cathemeral, but neither pattern was supported by PP values >0.67.

On both SF and LF reconstructions, the rates of transition from cathemeral activity to either nocturnal or diurnal imply that the diurnal and nocturnal niches may be more favourable for mammals. However, our results unequivocally support the persistence of cathemerality in mammals since the K-Pg. In primates, cathemerality has been argued adaptive under fluctuating environmental conditions^23,41^ and cathemeral species show higher speciation rates (although lower overall diversification rates) compared to nocturnal and diurnal species^24^. If these patterns are also true for the rest of Mammalia, they could explain the persistence of mammal cathemerality against the net outflow of species and slow diversification rates. In Lepidoptera (moths and butterflies), the persistence of a mixed (cathemeral) diel activity pattern has been argued to be the result of conflicting predation pressures, from bats during the night and birds during the day^42^. Hence, cathemeral activity may be preferred when strong selective forces are acting at opposite directions. The appearance of mammal cathemerality may have been due to high nocturnal predation risk, perhaps from other mammals that would have made the nocturnal niche less advantageous.

The higher transition rates for the LF tree are likely a result of the method we used to construct the species-level LF phylogeny, i.e. re-scaling the branch lengths of species-level clades from the SF phylogeny^28^ to maintain the length of the corresponding terminal branch provided by^27^. SF branch lengths were usually scaled down in this process, because the SF generally estimates older divergence dates than the LF, reflecting the difference between the two phylogenetic models. A consequence of our grafting procedure is that a band of artificially short branches is formed near these graft points, which implies rapid change. Higher rates allow for more change along tree branches, and reduce the precision of the results, which probably contributed to our LF reconstruction yielding fewer decisive predictions and lower statistical support compared with the SF reconstruction (Figs. 2, 3 and 4). Whilst a direct comparison of transition rates between the two phylogenetic hypotheses is therefore precluded, the broad pattern of transitions (i.e. low transition rates into cathemerality and high transition rates out of it in either direction) is supported in both analyses, as is the general pattern of temporal niche evolution that emerges from the node reconstructions.

Although we have demonstrated the importance of the phylogenetic comparative approach to the investigation of the evolution of behavioural traits in mammals, ancestral reconstruction methods rely heavily on the accuracy of phylogenetic estimates. The LF hypothesis of mammalian evolutionary history is well supported^27,37,40^, but phylogenetic estimates are only available at family-level, and further modification was required to add the species-level information for our analysis. Despite the attention attracted recently by studies of mammalian phylogenies^27,37,40,43^, only the SF hypothesis is represented by a species-level phylogeny, making the incorporation of the LF hypothesis and the explosive model problematic for phylogenetic comparative analyses that are based on detailed species-level data.

In conclusion, we argue that the activity patterns of Mesozoic mammals are consistent with the prediction of temporal partitioning, and that the gradual acquisition of daytime activity in mammals, first cathemerality then diurnality, coincided with the decrease in pressure from dinosaurs, whether due to their decline or extinction. Given the current evidence, temporal partitioning within Mesozoic amniotes mostly followed the phylogenetic (mammal-archosaur) division, but while some dinosaurs invaded the nocturnal niche^44^, we find little support for Mesozoic mammals invading the diurnal niche. The constraints on mammals becoming diurnal during the Mesozoic would have been strong enough to counteract the ecological pressure to diversify, following at least 100Myr of mammalian sensory and eco-morphological radiations that sub-divided their nocturnal niches. Mammals diversified rapidly once they expanded outside the nocturnal niche, but whether invading the diurnal niche facilitated mammals’ Cenozoic success remains to be answered.

## Methods

## Data

We collated activity records for 2415 mammal species, representing all 29 extant orders and 135 of 148 extant families from the PanTHERIA database^1^, and from published sources such as research articles, field guides, and encyclopaedias (Supplementary Table 1). Data collection was designed to incorporate extensive phylogenetic diversity, seeking to represent maximum ordinal diversity before targeting familial and generic diversity. We assigned each species into one of five activity patterns: (*i*) nocturnal - active only or mostly in the dark; (*ii*) diurnal – active only or mostly during daylight hours; (*iii*) cathemeral – active both during the day and during the night; (*iv*) crepuscular – active only at twilight, around sunrise and/or sunset; and (*v*) ultradian – active in cycles of a few hours. We follow the taxonomy and species binomials in Mammal Species of the World, 3^rd^ Edition^45^, with one exception: we use Cetartiodactyla, instead of separate orders Artiodactyla and Cetacea, following^46,47^. We resolved conflicts where sources disagreed on species activity pattern as follows: (i) records with a combination of either nocturnal and crepuscular, or diurnal and crepuscular were changed to nocturnal or diurnal, respectively; (ii) records from complied sources were preferred over localised studies (which are prone to idiosyncrasies); and (iii) records from more recent sources were preferred. This left 29 species unresolved and these species were excluded from subsequent analyses.

## Phylogenetic framework

We used two phylogenetic frameworks representing two of the main hypotheses of mammalian evolutionary history for our analyses: the short-fuse (SF) hypothesis is represented by the species-level “best dates” supertree^28^ updated from^29^, and the long-fuse (LF) hypothesis is represented by the amino-acid supermatrix phylogeny^27^ (Fig.1). The SF hypothesis asserts that the most recent common ancestor (MRCA) of all extant mammals diverged into its daughter lineages (Prototheria and Theria) in the mid-Jurassic, 166.2Mya, whereas according to the LF hypothesis this divergence took place in the late-Triassic, 217.8Mya. Both hypotheses agree that multiple extant lineages diverged in the Cretaceous and survived the K-Pg event (Fig. 1), but the SF hypothesis posits that intra-ordinal divergence of placental mammals had already begun prior to the K-Pg event, while the LF hypothesis places intra-ordinal divergence in the Cenozoic. A third evolutionary hypothesis, the explosive model, is supported by fossil evidence and morphological data^43^, but has been criticised for implying impossibly-high rates of evolution in the early-Cenozoic radiation of placental mammals, and for other problems^37,48,^ so we do not consider it here.

Here, we represent the LF hypothesis using the family-level supermatrix phylogeny^27^ (downloaded from TreeBASE: http://purl.org/phylo/treebase/phylows/study/TB2:S11872 on 01MAR2015). For our analyses we rendered it ultrametric, i.e. all the tips (species) of the tree are equidistant from the root, so that branch lengths are proportional to time. The LF hypothesis has recently gained support from several studies^37-40^, but it lacks species-level resolution, which is essential for our analysis. We therefore used each terminal branch of the supermatrix phylogeny (representing a taxonomic family) as a root branch onto which we appended the internal branching pattern of the family, as given in^28^ updated from^29^. In order to retain the original LF timeline, we scaled the appended branching pattern to 85% of its original supermatrix phylogeny branch length, and the root branch completed the remaining 15%. For this process we used functions from packages *ape*^49^ and *phangorn^50^* in R version 3.2.3^51^. Species that we had data for but that were absent from the phylogenetic frameworks were omitted from the analyses: 33 species from the SF phylogeny, and an additional 38 species missing from the LF phylogeny as families Aotidae, Pitheciidae and Lepilemuridae (Primates) were not originally included in the supermatrix phylogeny^27^. Thus, our analyses consist of 60% nocturnal species (n = 1399 species; n = 1384 species for SF and LF phylogenies, respectively), 14% cathemeral species (n = 321species SF; n = 320 species LF), and 26% diurnal species (n = 610 species SF; n = 588 species LF).

### Analyses

We used *BayesTraits υ3*^30^ to reconstruct the evolution of mammalian activity patterns. *BayesTraits* implements Markov Chain Monte Carlo (MCMC) methods to sample from the posterior distributions of transition rates for a transition matrix describing the evolution of a discrete character. The obtained posterior distribution allows the user to infer the posterior probability of each character state at the root and at each internal node of the phylogeny. By employing reversible jump MCMC (rjMCMC), *BayesTraits* is also able to sample from the posterior distribution of model configurations and optimise the number of parameters in the model. This removes the need for comparing models with different number of parameters by sampling from model space and parameter space concurrently^52^. We only consider the three main activity patterns across mammals in our analysis (nocturnal n = 1932 species, diurnal n = 999, and cathemeral n = 415, Supplementary Table 1) in order to reduce the complexity of the model and increase its biological interpretability (four transition rates instead of 16). Additionally, we do not consider ultradian activity patterns as these are mostly found with polar and subterranean species, where the 24-hour cycle is of reduced importance. We consider an ordered model of trait evolution: Nocturnal↔Cathemeral↔Diumal, whereby direct Noctumal↔Diurnal transitions are not allowed (set to zero), because morphological and histological adaptations to diurnality and nocturnality are mutually exclusive, while cathemerality involves an intermediate state of the relevant phenotypes^20,53^. Our underlying hypothesis is that during shifts from diurnality to nocturnality (or vice versa) species go through a phase where they are equally well adapted to both. All other transition rates were free to take any value. We used rjMCMC to estimate the optimal model configuration^52^. As activity pattern in our analyses was not a binary trait, we used the ‘multistate’ mode of *BayesTraits* to sample from the posterior distribution of transition rates between activity pattern categories. For each phylogeny, we opted for the reversible-jump MCMC procedure, and set a wide uniform prior, bounded between 0 and 100 for all transition rates, to ensure that our prior did not have a strong effect on the nature of the posterior. Each rjMCMC chain was run until convergence was reached (at least one million iterations), after which point the chains were sampled every 4000 iterations until a posterior of 1000 samples was obtained. We chose this wide sampling interval in order to minimise autocorrelation in our posterior samples. We ran twelve replicates of each chain (corresponding a phylogeny) in order to ensure consistency, and that each independent run converged on the same posterior distribution. The marginal likelihoods of each chain were calculated using the stepping stone sampler^54^ as implemented in *BayesTraits* (500 stones, 1000 iterations per stone) and compared between independent replicates to ensure consistency. In order to estimate the character state at each internal node, we used the modal value of the PP of each character state, calculated as the peak value of the kernel density of each posterior distribution. We used the R package *phytools* ^55^ to plot the PP values of each node on the mammal phylogenies (Figs. 3 and 4). To measure the accumulation of mammalian temporal niches over time, we calculated the running total of nodes (lineages) where an activity pattern was supported with PP > 0.67, and plotted this along the mammal evolution timeline (Figs. 3 and 4). The confidence threshold of 0.67 was chosen because an activity pattern supported at this level would mean that the next most likely pattern is less than half as likely.

### Data Availability

The authors declare that all data supporting the findings of this study are available within the paper and its supplementary information files. All data have been deposited on Figshare and will be made publically available after manuscript acceptance (doi: <u>10.6084/m9.figshare.4775416;</u> doi:10.6084/m9.figshare.4774648). Reprints and permissions information are available at www.nature.com/reprints.

### Code Availability

Computer code essential for replicating the results in this study has been deposited on Figshare and will be made publicly available after manuscript acceptance (doi: 10.6084/m9.figshare.4797367).

**Supplementary Information** is available in the online version of the paper.

## Acknowledgements

We thank TCD Lucas, S Meiri, EE Dyer, O Comay and I Pizer-Mason for technical assistance and providing data, and N Kronfeld-Schor for discussion. This work was funded with support from ISF grant No. 785/09 (TD), Tel Aviv University GRTF fund and Naomi Kadar Foundation (RM), and NERC Open CASE PhD studentship (NE/H018565/1) (HFG).

## Author Contributions

RM, TD and KEJ developed the overall study design. RM collected and processed the data, and carried out the analyses with assistance from HFG. RM and KEJ led on the writing of the manuscript with significant contributions from all authors.

## Author Information

The authors declare no competing financial interests. Correspondence and requests for materials should be addressed to RM or KEJ.

